# Can grid cell ensembles represent multiple spaces?

**DOI:** 10.1101/527192

**Authors:** Davide Spalla, Alexis Dubreuil, Sophie Rosay, Remi Monasson, Alessandro Treves

## Abstract

The way grid cells represent space in the rodent brain has been a striking discovery, with theoretical implications still unclear. Differently from hippocampal place cells, which are known to encode multiple, environment-dependent spatial maps, grid cells have been widely believed to encode space through a single low dimensional manifold, in which coactivity relations between different neurons are preserved when the environment is changed. Does it have to be so? Here, we compute – using two alternative mathematical models – the storage capacity of a population of grid-like units, embedded in a continuous attractor neural network, for multiple spatial maps. We show that distinct representations of multiple environments can coexist, as existing models for grid cells have the potential to express several sets of hexagonal grid patterns, challenging the view of a universal grid map. This suggests that a population of grid cells can encode multiple non-congruent metric relationships, a feature that could in principle allow a grid-like code to represent environments with a variety of different geometries and possibly conceptual and cognitive spaces, which may be expected to entail such context-dependent metric relationships.

## I. INTRODUCTION

Grid cells appear to comprise an essential component of the cognitive representation of space in rodents [1] and in other species, e.g. bats [2]. Located in the medial entorhinal cortex, these neurons are selectively active when the animal is in certain positions of the environment, the so-called *fields*, at the vertices of a remarkably regular hexagonal lattice. A study of the activity of grid cells in multiple environments [3] has shown that while the grid expressed by each neuron varies across environments in its spatial phase and orientation, between neurons the coactivity relations are largely preserved, at least for those recorded nearby in the tissue, with the same tetrode. In other words, the grids of different cells undergo a coherent rigid movement when a new environment is explored, as illustrated schematically in Fig.1 (a) and (b). The subsequent discovery of quasi-discrete “modules” [4] indicates that these relations are maintained at the local network level, presumably by recurrent collateral connections among grid cells. This finding has led to the hypothesis that local ensembles of grid cells comprise each a single continuous attractor network, expressing a “universal”, two-dimensional map, which encodes the metric of space independently of the environmental context. There is a crucial difference with the context-dependent spatial representations provided by hippocampal place cells, which display “global remapping” [5] even between very similar rooms, in particular in the CA3 field [6]: cells which were silent acquire one or more place fields, others lose theirs, and the fields that seem to have been maintained typically are in a different location (Fig.1B). Global remapping has motivated the conceptual model of multiple charts [7], in contrast with early and later models of continuous attractor grid cell networks, which envisage a single chart [8],[9],[10]. The dominant overall view, then, holds that the hippocampus encodes multiple, uncorrelated, context-dependent cognitive maps, while the grid system provides metric information that is independent of the environment. Recent evidence of context-dependent distortions in the grid pattern have begun to question the view that the collective map expressed by a grid module is universal, that is, that it applies to any environment. Stensola et al. [11] have shown that, when rats explore large environments, a single grid can exhibit multiple orientations, likely due to anchoring effects to the closest wall, which in any case amount to distortions of the hexagonal pattern. These effects have been analyzed extensively in a more recent study [12]. Krupic et al. [13], [14] have shown that the grid pattern deviates from perfect hexagonality, with both global and local distortions, in response to environmental features such as the geometry of the walls. Finally, a couple of recent studies [15],[16] have shown that the presence of salient features such as goals or rewards affect the entorhinal map, changing field locations and inducing remapping in other space selective cells. These observations, moreover, refer solely to the position of the peaks of activity, i.e. the place fields of each cell, and do not take into account the fact that they vary reliably in height, independently across peaks, from one environment to the other [17]. Should we still regard grid cells as a sort of stack of millimeter paper, providing a universal metric for space?

**FIG. 1.**
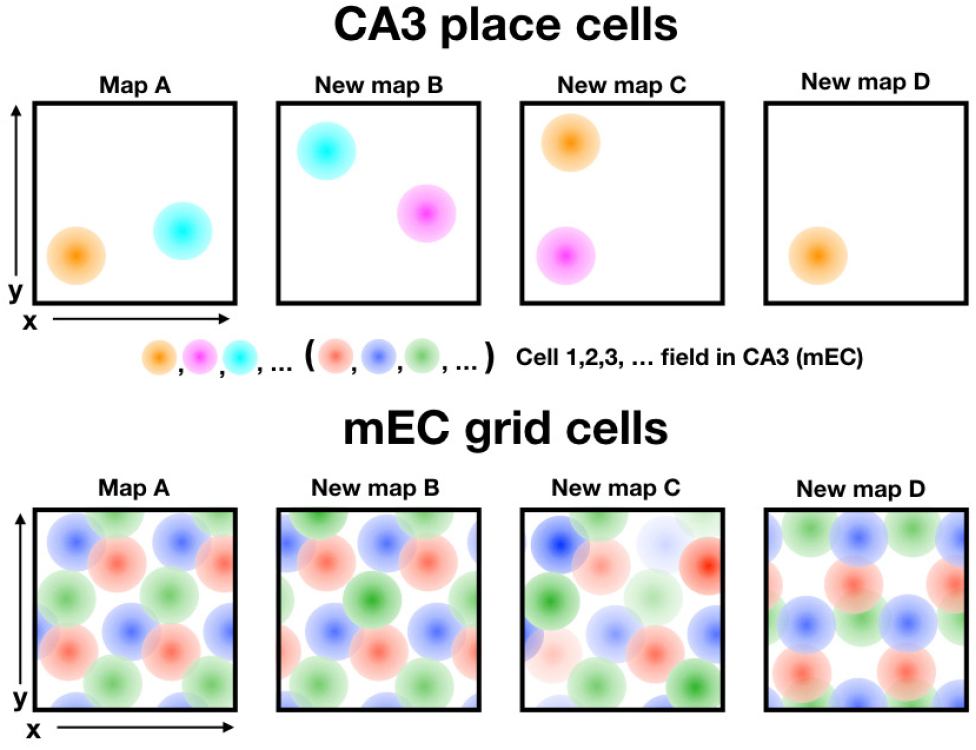
Types of change in grid cell activity in mEC (bottom) concurrent with global remapping in the CA3 field of the hippocampus (top). The universal grid map model, idealized from [3] allows only for a coherent translation (and possibly a rotation) into a new map B, when changing environment. Under a manipulation which does not entail changing environment, the individual fields of each unit have been observed to independently vary their peak rates, keeping their relative position ([20], new map C). We assess the hypothesis that the same network may also express other maps, such as map D, with a complete re-positioning of the grids of different units.

Recent studies conducted in both rodents and humans, moreover, suggest that regular grids may not “measure” only physical space. Aronov and colleagues [18] find that both place cells and grid cells, in rats, are involved in the representation of a non-spatial but continuous, one-dimensional variable, such as the frequency of a sound. An fMRI study by Constantinescu et al. [19] shows an hexagonal modulation of the BOLD signal in human Entorhinal Cortex, and elsewhere, in a task that requires subjects to “navigate” the 2D space spanned by the varying leg and neck lengths of a drawing of a bird. The representation of abstract or conceptual spaces, which could in principle be topologically and geometrically complex, would require of the grid cell system a flexibility that can hardly be reconciled with the universal grid hypothesis.

In a most interesting study [20], a subset of grid units were depolarized in transgenic mice, leading to what appears to be global remapping in the hippocampus. What is so striking is that the manipulation induces extensive changes, up and down, in the peak firing rates of the different fields of individual grid units, but not in their position. This elaborates the observation in [3], and suggests that what might be universal in the grid representation expressed by an ensemble of units, if anything, are the relative positions of the fields, whereas their peak firing rates are variable (Fig.1C). On the other hand, a strict hexagonal periodicity of the field positions of individual units is only possible in flat 2D environments. The adaptation model of grid formation [21] predicts instead, on surfaces with constant positive or negative Gaussian curvature, and appropriate radius, the emergence of grids with e.g. pentagonal [22] or heptagonal [23] symmetry. In all other cases, including ecologically plausible natural environments, non-flat surfaces have varying curvature, making strictly periodic grids dubious, and rigid phase coherence most unlikely. But then, what happens to the universality of the grid in natural environments?

To address these issues, the aim of the present work is to answer a first fundamental question: is it at all possible to conceive of multiple, hence non-universal, ideal grid representations expressed in the same local network, when the animal is placed in *distinct*, even if flat, environments? In other words, would the storage capacity of a recurrent network of grid cells be above unity, so that multiple continuous attractors can coexist, encoded in the same synaptic efficacies? We pose this question within two alternative mathematical models, both accepting the idealized assumptions which underlie the universal map hypothesis, that is, of strict periodicity and equal peak rates, depicted in Fig.1D, but allowing for several uncorrelated grid representations. Under these assumptions, we analyze an ensemble of grid cells as a Continuous Attractor Neural Network, extending the frameworks developed in [24], [25] and [26] for the description of place cells. We emphasize that the storage capacity we are interested in quantifies the number of different, independent *charts* (or collective maps) that the network can store, and not the spatial resolution (which may be referred to as *information capacity*, i.e. the number of different positions that can be decoded from the ensemble activity), as considered for example in [27] and [28].

## II. COMPLEMENTARY NETWORK MODELS

We model the grid cell population as an ensemble of units interacting through recurrent connections, whose structure defines which activity states are robust - the dynamical attractors. We assume, however, that a separate process, based *e.g.* on adaptation [21], has determined the emergence of a periodic grid, independently for each unit, during familiarization with each of *p* distinct environments; meanwhile, recurrent connections are shaped by a Hebbian learning process, such that neurons that happen to have nearby fields tend to fire together, strengthening their connections, while neurons with fields far apart remain weakly connected. The connection strength *J*_*ij*_ is therefore taken to be a sum of contributions from the exploration of *p* environments, with each contribution, once averaged across many trajectories, a function of the relative position of the fields in that environment. Exploiting the simplifying assumption that each grid is strictly periodic, we can focus on the elementary repetitive tile, which has only one field per unit and is, in the mathematical formulation, connected by “periodic boundary conditions” to adjacent tiles. The assumption of periodic boundary conditions is motivated by the remarkable regularity of the arrangement of the fields observed in the original experiments, and by the model being meant to describe interactions within a grid module, in which all cells share the same spacing and orientation. The contribution to the connection strength between two units *i* and *j* is then reduced to a function of their field centers 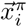 and 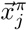 on the elementary tile in environment *π*

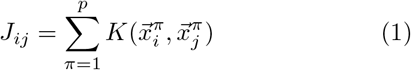

where we refer to *K*(⋅) as the “interaction kernel”. The field peaks, or centers 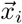 of *N* units are taken to be randomly and uniformly distributed over the elementary tile. Our analysis focuses on two different models of neurons (binary and threshold-linear) and two types of attractor symmetry (square and hexagonal), which stem from the tile shape or the interaction kernel. Both neuron models allow, from complementary angles, a full statistical analysis, leading to otherwise inaccessible results. The storage capacity turns out to depend more on how interference reverberates through loops (expressed by the parameter *ψ*, see below) than on the type of units; and interference, in the densely coded and densely connected regime, affects square much more than hexagonal grids.

### A. Binary units

The first model we consider is an extension of the model proposed by Monasson & Rosay [25] for the modeling of place cells in CA3. Here the activity of neurons is described by binary variables, such that the pattern of activity of a network of *N* units is a vertex {*σ*} ∈ {0, 1}^*N*^. For the binary model, the kernel *K*(*i, j*) between units *i* and *j* relative to one environment is taken to be a step function of the distance between their field centers

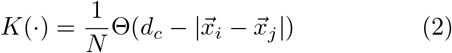

where Θ(*x*)=1 for *x >* 0 and 0 otherwise – note that the distance 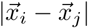 is along the shortest path, considering the periodic boundary conditions. The periodic structure of the attractor depends on the shape of the rhomboid unitary tile in which the field center 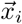 of each unit is located. The lattice symmetry is specified by the angle *θ* between its two primitive vectors. *θ* = 60° corresponds to the standard case of hexagonal grids, while *θ* = 90° describes a square grid pattern. These two cases and the resulting interaction kernel are depicted in Fig.2 (a) and (b). The cut-off distance *d*_*c*_ sets the number of non-zero connections each unit receives from the storage of a given environment, denoted by *wN*: 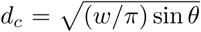. This measure of connectivity within one environment should not be confused with the global connectivity taking into account all environments, *C* = (*N* − 1)(1 − (1 − *w*)^*p*^) ~ *N* for large *p*.

**FIG. 2.**
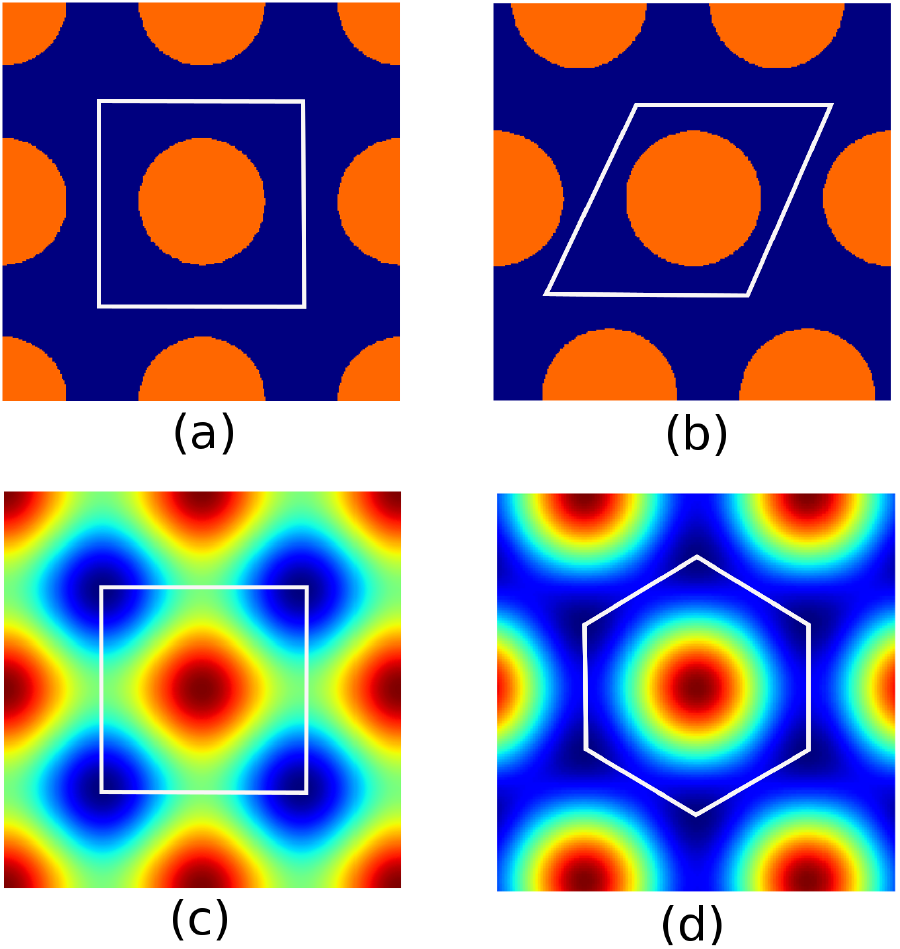
Interaction kernels for the binary (a,b) and rate (c,d) models. The white lines show the elementary tile of each lattice.

The dynamics of the network is governed by the energy function:

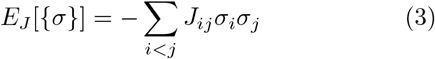

and constrained by the requirement that at any time a fixed fraction *f* of units be in the active state, i.e. ∑_*i*_ *σ*_*i*_ = *fN*. We call *f* the coding level, or sparsity of the representation. This constraint is taken to reflect some form of global inhibition. Later we shall focus only, given *w*, on the optimal coding level in terms of storage capacity, hence on a specific value *f*\*(*w*), which turns out to be a monotonic function of *w* (see Fig.3). This model then allows an explicit focus on the dependence of the storage capacity on the width of the kernel and on the resulting optimal sparsity of the representation.

**FIG. 3.**
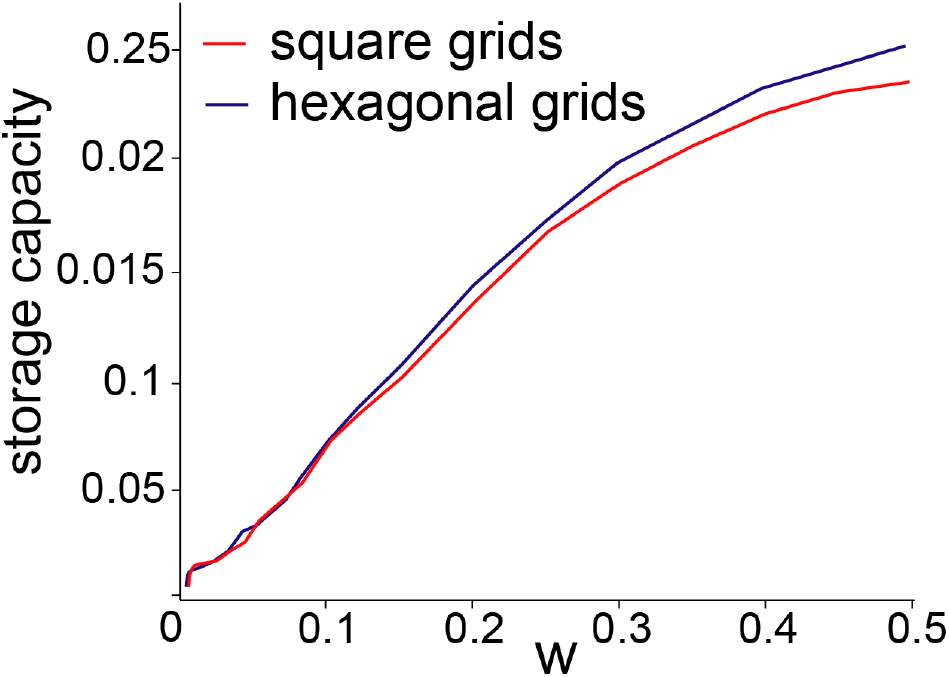
Optimal coding level for the binary model.

### B. Threshold-linear units

We extend our analysis to firing-rate units, whose activity is described by a continuous positive value corresponding to their instantaneous firing rate. This second model allows us to capture the graded nature of neural activity, which is salient when it represents space, which is itself continuous. The activity of the network is given by a configuration {*V*_*i*_} ∈(ℝ^+^)^*N*^, and each unit integrates the inputs it receives through a threshold-linear transfer function [29]

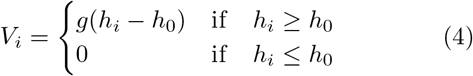

where *g* (the linear gain) and *h*_0_ (the activation threshold) are global parameters of the network, and the “local field” *h*_*i*_ is a real-valued variable summarizing the input influence on unit *i* from the rest of the network, which we take to come from a random but fixed set of *C* among the *N* − 1 other units, as well as from external sources. The interaction kernel *K*(⋅) is given by the special sumof-cosines form

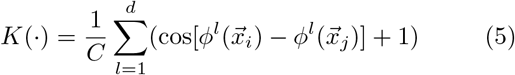

which had been considered as a toy case by [24], before the discovery of grid cells. The field center of each unit on the elementary tile is expressed by a set of angles 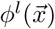. We shall see that *d* = 2 and 3 are equally valid choices on the plane, as well as *d* = 1, which leads to “band” solutions (see below). This model therefore allows decoupling the form of the kernel, which is extended, with interactions among units far away on the elementary tile (and the resulting coding level is correspondingly non sparse) from the connectivity, which can be made arbitrarily sparse if *C/N* →0. As a superposition of *d* cosine functions, the kernel can also be conveniently written as a sum of dot products. The +1 term is added to enforce excitatory connections. While not circularly symmetric like the radial kernel used in the binary model, this cosine kernel allows for the analytical study of periodic patterns that are spread out on the plane, with a large fraction of the units active at any given time. The solutions for the hexagonal kernel (Fig.2(d)), in particular, given by three cosine functions at a 60° angle from one another, have been considered as a reasonable model for experimentally observed grid cells. In the figure, the hexagonal elementary tile extends in the range *x* = *±*1*/*2 and 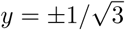, and the three angles span the directions 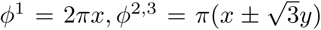. The square kernel is obtained for *d* = 2 and the two cosines at 90° from each other (Fig.2 (c)). Note that, as with the binary model, *N* units are concentrated on an elementary tile that in the hexagonal case is 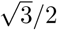 of the area of the square case.

An energy function would look similar to the one in Eq. [3], but now expressed in terms of the continuous variables {*V*}. When *C < N* − 1 and connections are not symmetric, the energy formalism does not apply but we can still analyze the model (see below and in appendix B), and again we take global inhibition, which can now also act through a modulation of the common gain *g*, to depend on the average activity of the network and to be such as to optimize storage capacity.

## III. STORAGE CAPACITY

Both models can store a single population map, as in the bottom panels of Fig.1A,B, and the equations for such a map admit periodic bump solutions that reproduce the shape of the tile/kernel (as well as potentially other solutions, e.g. stripes, to be discussed later). We are interested however in their capacity to store several distinct maps, as in Fig.1A and D, and in the possibility to calculate such storage capacity analytically, in the mean field approximation. The general strategy involves formulating and resolving a set of self consistent equations relating the activity of the units in the network. When the model admits an energy function, these are the saddle point equations derived from the computation of the “free energy” of the system with the replica trick, which allows to take into account the statistics of the field centers in each environment. Without an energy function, e.g. when the connections are sparse and not symmetric, equivalent equations can be derived through the so-called Self Consistent Signal-to-Noise Analysis [30]. The solutions to these equations, that describe the activity in one map, disappear sharply at a critical value *α*_*c*_ of the storage load *α* = (*p/C*), which measures the ratio of the number of maps to the number of connections to each unit. *α*_*c*_ then gives the maximum number of maps that the network can store and retrieve or express, normalized by the connectivity. Crucially, we have developed a novel method to assess whether below *α*_*c*_ these solutions are indeed stable and prevail on others (Fig. 6 and 7).

The details of these methods, that build on [25] and [26] for the binary model and on [31] and [24] for the rate model, can be found in appendix. We focus, in the calculation of the storage capacity, on so-called “bump states”, in which activity is localized along each of the two dimensions of the elementary tile (anywhere on the tile, given the translation invariance of the interaction kernel). Other solutions however exist, as discussed in section IV.

### A. Binary units

The statistical analysis of the minima of the free energy leads to the patterns of activity {*σ*} that are likely to be observed given the connectivity. More precisely, we have derived self-consistent equations for the average activity 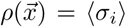 of unit *i* having its grid field centered in 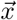 (in the elementary tile):

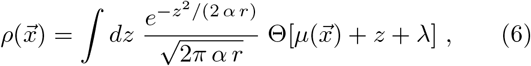

where

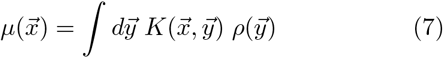

is the signal input received by the unit through the interactions corresponding to the environment in which the bump is localized, say, *π* = 1, and *z* is the noisy, Gaussian input due to the interference from the other environments, say, *π* = 2*,…, p*, see Eq. (1). The variance *α r* of these Gaussian inputs is, in turn, self consistently derived from the knowledge of the activity profile *ρ*, see appendix A. The uniform (inhibitory) input *λ* enforces the constraint 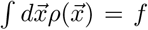. We have considered the limit case of neurons responding deterministically to their inputs, although the analysis extends naturally to stochastic noise.

We calculate, from the saddle point equations, the storage capacity *α*_*c*_(*w, f*) as the maximal value of the load *α* for which a bump-like solution to Eq. [6] exists. Then, for a given value of *w*, we find the coding level *f*\*(*w*) that maximizes the storage capacity. Over a broad range 0 ≤ *w* ≤ 0.5 the optimal *f*\* turns out to be approximately half the value of *w* (see Fig.3). That the optimal value for the coding level is proportional to *w* can be understood intuitively by considering the spatial profile of the signal 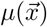: if too few cells are allowed to be active, the connections to the cells that are forced to be silent, within the connectivity range of the active cells, will be frustrated. On the other hand, if too many cells are active, those outside the connectivity range will contribute more to the noise than to the signal. This optimal storage capacity is plotted in Fig.4, for the square and hexagonal grids as a function of *w*. At low *w* the two values are similar, but when *w* increases their trends diverge – a *ψ*-related effect – leading to substantially higher capacity value in the hexagonal case, of order 10^*−*2^ for *w* ≃ 0.5. This value would definitely allow, in a real cortical network with order thousands (or tens of thousands) of neurons, the storage and retrieval of multiple independent grid maps. Again considering the spatial profiles of the signal 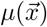 allows to gain intuition about this divergence. At very low *w*, i.e. short range interactions, what happens in other tiles can be neglected, and the two grids behave similarly. When the range is wider, the location of the fields in the immediately neighbouring tiles starts to be relevant. In the square case, there are four first neighbours, contributing to excite silent neurons in-between the fields. For an hexagonal arrangement of the fields, there are six neighbouring tiles that each contribute relatively less excitation in-between fields. Intuitively this last geometrical arrangement makes the structure more rigid and reduces the influence of the noise due to the storage of other charts.

**FIG. 4.**
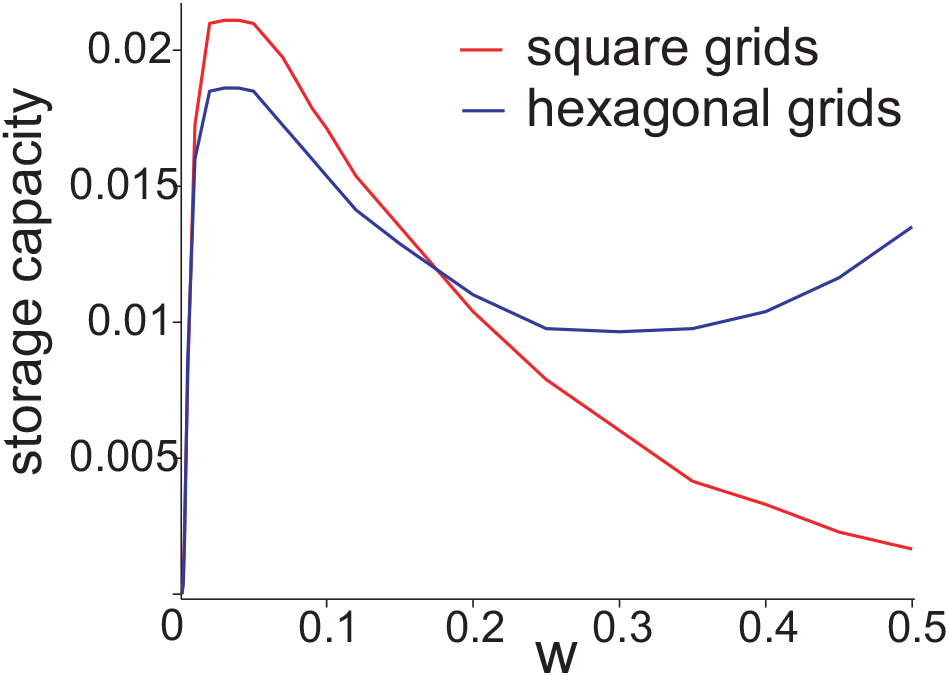
Storage capacity as a function of w for square and hexagonal grids in the binary model, given an optimal coding level f ≃ w/2.

### B. Threshold-linear units

In this model the coding level and the connectivity range are both fixed by the shape of *K*(⋅). The mean field approach can be however extended to the case of arbitrary values of the connectivity density *C/N*, with the Self-Consistent Signal-to-Noise Analysis [30]. The storage capacity is given by the *α* for which the solution to the equation

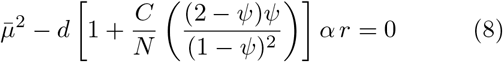

disappear. In fact, the disappearance of the solution only gives an upper bound on *α*_*c*_, as one has to check its stability as well. The details of the derivation and the expression of the average signal 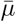 and of the interference noise *r* are reported in appendix B. We plot such critical value for square and hexagonal grids with the respective kernels, as a function of the inverse density *N/C*, in Fig.5 (full curves, blue and red). In the fully connected regime, we find a result, corroborated also by computer simulations, similar to the one obtained with the binary model, with however a huge difference in capacity between square and hexagonal grids, and a value ~10^*−*2^ only for the latter. Moreover, it turns out that for the square kernel the stripe or band solutions of the next section are the global minima, and the square solutions are only marginally stable. In all cases the capacity increases as the connectivity density decreases, reaching an asymptotic value as *N/C* → ∞. The quantitative results for hexagonal grids has implications consistent with those of the binary model: it suggests that, again, a network of grid cells, for which a plausible number of synapses per neuron may be in the order of thousands, and with a connectivity, say, of order *C/N* ≃ 0.1, would have the capacity to encode perhaps a hundred different environments.

**FIG. 5.**
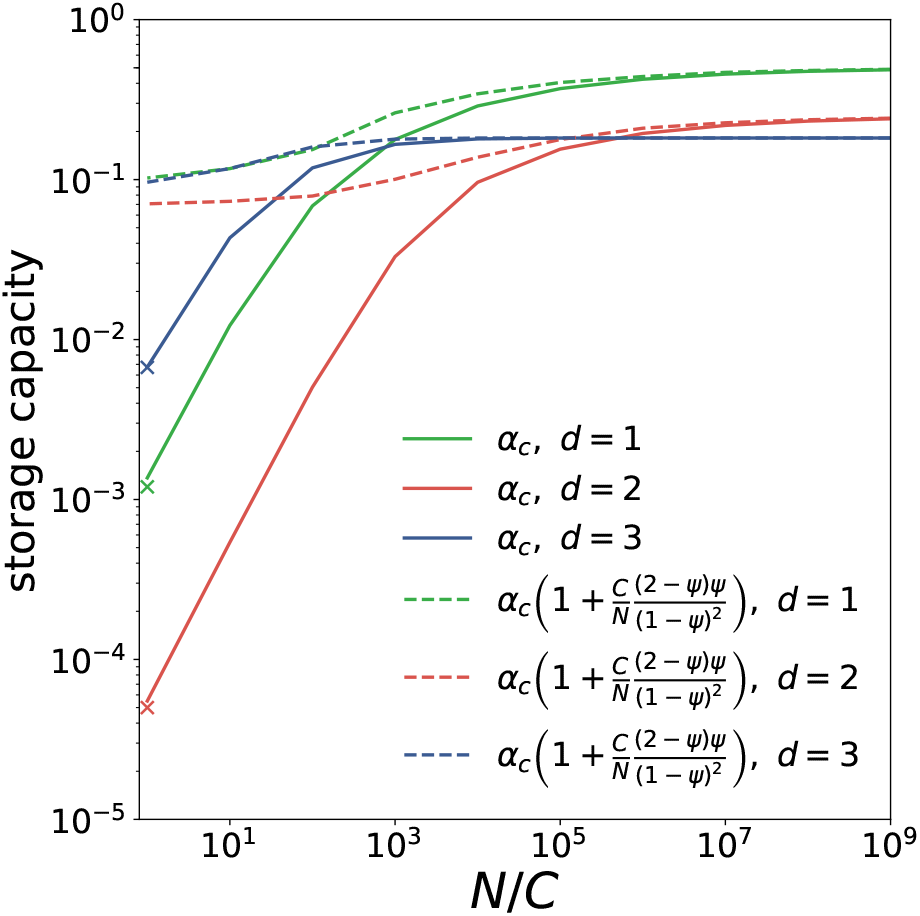
Storage capacity in the threshold-linear model as a function of the inverse connectivity density N/C, on a log-log scale. Full lines give α_c_ for the three different interaction kernels (bands in green, square grids in red and hexagonal grids in blue). Dashed lines indicate what the capacity would be without noise reverberation. The crosses on the left show the capacity obtained with numerical simulations for a fully connected network.

### C. Sparsity and noise reverberation

The binary model shows that the difference in capacity between hexagonal and square grids results from the effective interactions among the fields in different tiles, as it emerges only with wide kernels and dense coding. When both are sparse, hexagonal and square grids are roughly equivalent. The *w →* 0 limit can be worked out analytically and *α_c_ →* 0 in both cases, but only after having reached a maximum around *α*_*c*_ ≃ 0.02 for quite sparse codes, *w* ≃ 0.03 and *f* ≃ 0.015. Sparse coding is known to suppress noise reverberation (leading to small *ψ*), but remarkably this relatively large capacity is approximately preserved for hexagonal grids with dense coding, *w* ≃ 0.5 and *f* ≃ 0.25, illustrating the efficiency with which this compact arrangement minimizes interference.

The threshold-linear model affords complementary insight, again on how the hexagonal/square capacity difference depends on the units active in each attractor reverberating their activity. Mathematically, this is expressed explicitly by the dependence of Eq.8 on the order parameter *ψ*, which quantifies the amount of reverberation through the loops in the networks. The physical meaning of *ψ* can be inferred from the expression derived in appendix B and C:

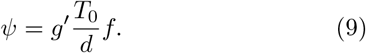

The factor *g*′*T*_0_*/d* is in fact the typical noise *T*_0_*/d* amplified by the renormalized gain *g*′ and multiplied by the average fraction of active units, the *f* parameter as in the binary model. *ψ* is then the one-step loop term in the reverberation of the noise; its effect on the capacity is illustrated by the dashed line in Fig.5, in which such contribution is factored out. For densely connected networks, storage capacity would massively increase and relative differences would decrease without noise reverberation. The optimal capacity for the hexagonal kernel is then (mainly) the result of a reduced reverberation of the noise, due to the shape of the activity distribution of its attractors: the average fraction of active units (*f* ~ 0.46) in the attractive state of the hexagonal kernel model is considerably lower than the same fraction in the square kernel, where it would be *f* ~ 0.79 for the square grids, and is only somewhat reduced to *f* ~ 0.68 for the stripes, which replace them as the stable solutions for this kernel.

## IV. BAND SOLUTIONS

In the previous analysis, we focused on “bump” states, in which activity is localized in a grid pattern. Another possibility are partially localized solutions: “band” states, where activity is localized along a single direction in the elementary tile, and extends along a stripe in the orthogonal direction.

In the binary model, these band states can be oriented along an edge of the tile (Fig.6(b,f)), or along the diagonal of the tile (Fig.6(c,g)), or in a discrete multiplicity of other orientations. Individual units “fire” along stripes of the same orientation, with relative offsets. We can study the propriety of some of these band states in the *w − f* parameter space, to find that they are particularly favored in regions of high coding level. Given the connectivity range set by *w*, bump states are the global minima of the free energy for low *f*, and one of the band states (which one depends on *θ*) becomes the minimum for higher *f*. For example, for both square and hexagonal grids, at small connectivity range *w* = 0.1, band states have lower free energy than the bump state for coding levels beyond 0.25, while for the larger connectivity range *w* = 0.5, this happens for coding levels beyond 0.4. This is intuitive, since for sufficiently large *f* a band state has a shorter boundary between active and quiescent units than a bump, and it is the length of the boundary that raises the free energy above its minimum value. Moreover, we can study how these different states are separated by computing the size of the free-energy barrier to cross to go from one state to another. The method to compute this barrier is sketched in Fig.7(c) and explained in more details in appendix D. In Fig.7(d) we show the size of the barriers to cross to go from a “bump” state to “band” states. On the range of coding levels where these two kinds of states co-exist, the “bump” state is always more robust for an hexagonal grid compare to a square grid, as shown by the higher barrier size in an hexagonal grid (blue curve, from Bump to Band Edge or Band Diag. state) compare to square grid (full red curve, from Bump to Band Edge state).

A different behaviour is observed in the threshold-linear network. In this case, the rigid symmetry imposed by the 3-cosine interaction kernel makes the bump pattern a global minimum. In the 2-cosine case, instead, band state are stable solutions, corresponding to a macroscopic overlap with only one of the two cosines. We can describe bands also with a 1D interaction kernel, with a single cosine, and compare the storage capacity for band patterns with the one for square and hexagonal grids. In Fig.5, the green line shows the capacity for band patterns as a function of the connectivity. For a densely connected network, it is above that for square grids, and the barrier methods indicates that these are only marginally stable to collapsing into stripes. This is in line with the reduction of the capacity from one to two dimensions shown in [24]. Interestingly, as soon as the emergence of a third cosine is allowed the capacity is instead enhanced, surpassing the 1D kernel except for very low values of connectivity density.

**FIG. 6.**
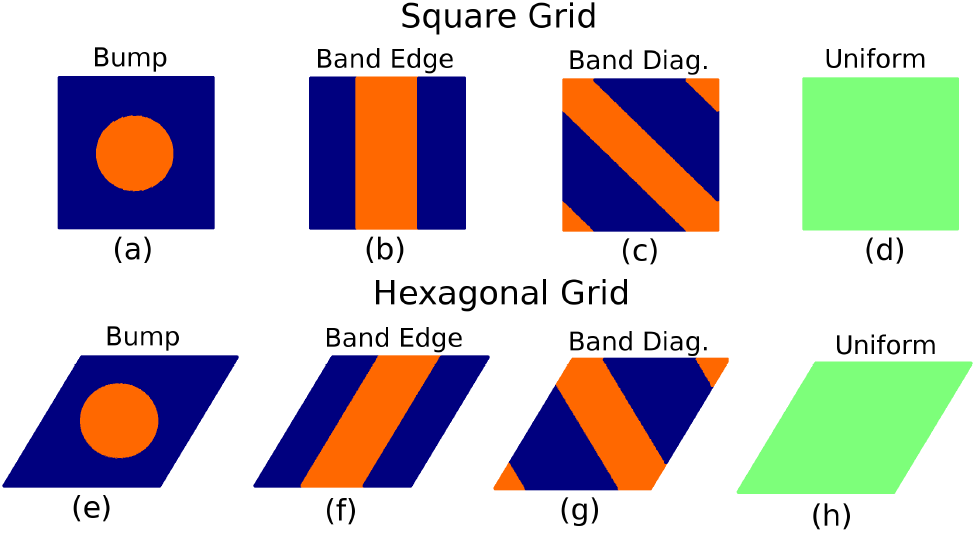
Different solutions to the saddle point equations in the binary model. Bumps (a,e) are stable at low f (f=0.2 in the figure). Edge-oriented and diagonal bands are stable solutions for the θ = 60° tile at higher f (e.g. f=0.4, f,g), but only the former (b) are stable for θ = 90°. Uniform solutions (d,h) are always unstable below the critical capacity.

**FIG. 7.**
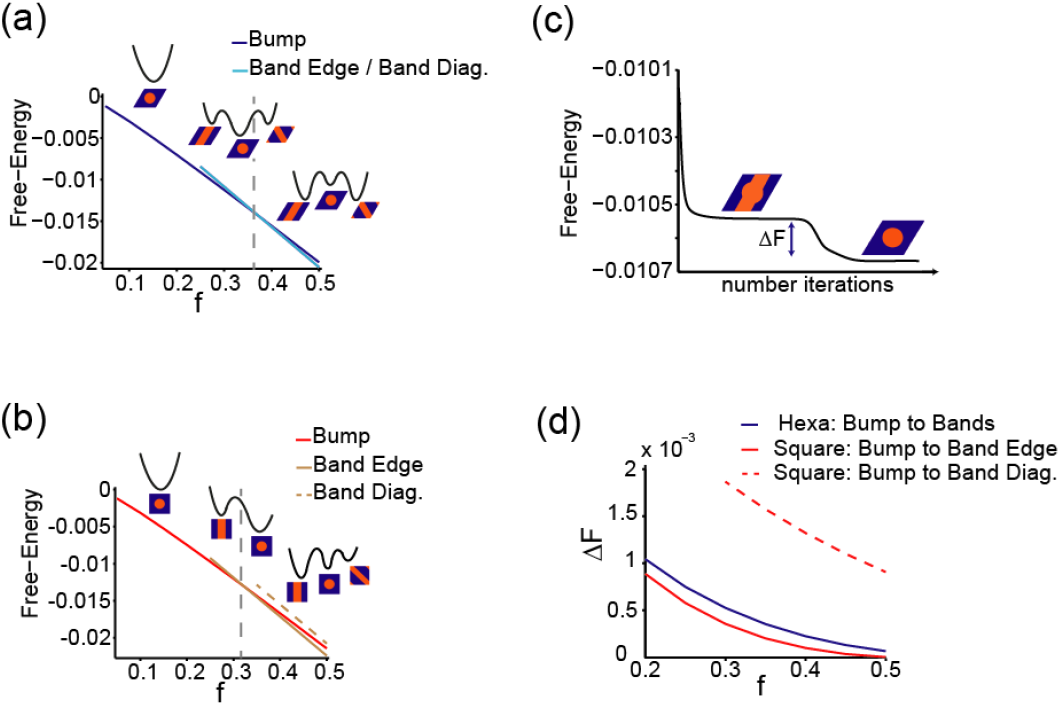
Bump and band states in the binary model. Free-energies of the bump and band states for hexagonal grids (a) and square grids (b). (c) Free-energy barriers are given by the difference in free-energies between an unstable mixed-state (band edge + bump shown here) and a metastable state (bump state shown here). (d) Size of the free-energy barriers to cross to go from the bump state to band states. w = 0.1, α → 0.

## V. DISCUSSION

Our results indicate that, given appropriate conditions, a neural population with recurrent connectivity can effectively store and retrieve many hexagonally periodic continuous attractors. This possibility suggests that a regular grid code may not be restricted to represent only physical space; it could also express continuous abstract relations between arbitrary features, at least if they can be mapped to a two-dimensional space. This would however require a system flexible enough to store and retrieve uncorrelated grid representations. Our results show that this flexibility does not need, in principle, separate neural populations for separate representations, but can be achieved by a single local ensemble, provided it can learn effectively orthogonal representations.

Given the recent observation of non-spatial coding – a consistently tuned response to the “position” along a 1D non-spatial variable, sound frequency, during a sound manipulation task – by neurons that qualify as grid cells in a 2D spatial exploration task [18], it would be interesting to know whether a similar selectivity can be observed for a 2D non-spatial variable, as suggested by indirect observations of hexagonal modulation [19]. Several important questions are left open for future investigation. First of all, if global remapping is possible within a grid cell population, why has it not been observed experimentally? Possibly, a remapping capacity of grid cells may have been hidden by the fact that multiple mappings were only studied in simple, empty, flat environments and then they turned out to be the same, modulo translations [3]. The hypothesis of a universal grid, that shifts without deformation across an environment and from one environment to the other, faces severe difficulties as soon as curvature is taken into consideration. In curved environments, rigid translations are not possible, and the geodesic transformations that partially substitute for them do not leave field-to-field relations unchanged, making a universal grid *a priori* impossible. Nevertheless, natural environments show a wide range of both positive and negative curvature, which does not seem to pose any problem to the navigational skills of rodents, or of other species. It is then conceivable that the apparent universality of the grid pattern comes from the experimental restriction to flat environments, which all belong to the same, rather special, class of two dimensional spaces with zero curvature, and that a richer grid behavior is required in order to code for position in more general spaces. The emergence of grid representations in curved environments has been investigated with a model based on single cell adaptation [22][23], which illustrates the emergence of different regular patterns for distinct ranges of curvature. Estimating the storage capacity of recurrent networks expressing curved grids, however, poses some challenges. Since shifting the grid pattern along a curved surface moves individual fields by a different amount, the relationships between grid units cannot be reduced to the relationships between a single pair of their fields. Long-range translational coherence becomes impossible. Curved grids can be only partially coherent, and whether this partial coherence is sufficient to build stable attractors is an open problem [32]. A second open problem is the ability of a network encoding multiple charts to support path integration, since the noise introduced by other charts is likely to introduce discontinuities in the dynamics shifting the activity bump, impacting the accuracy of the integrator. It has recently been suggested [33] that interactions between different grid modules (each encoding a single chart or coherent ensemble of maps) can enhance the robustness to noise during path integration. The possibility that this result generalizes to modules encoding multiple charts, and the analysis of the capacity deriving from interactions between modules, are beyond the scope of the present work, but deserve future investigation. Finally, a third issue concerns the learning dynamics that sculpts the grid attractors. What is the mechanism that leads to the attractors of the recurrent network? Does a single grid dominate it, in the case of flat environments? Can *self-organization* be unleashed by the interplay between the neural populations of mEC, including non-grid units, and hippocampal place cells, aided by the dentate gyrus [34]? Including the hippocampus may be needed also to understand the distortion of the grid pattern, reported in several experimental studies [4][11][13], that by disrupting long-range order also weakens coherence. At the system level, a finite storage capacity for the grid cell network implies the possibility that medial Entorhinal Cortex, or any other area in the brain [19] that is observed to include grid-like units, can serve context memory. This would turn upside down the widely shared notion that memory for the specific spatial features of each environment is only available downstream, in the hippocampus, and conceptually reunite medial Entorhinal Cortex with other regions of the mammalian temporal lobe, known to be dedicated to their own flavour of memory function [35].Moreover, the possibility of multiple uncorrelated continuous attractors in flat environments, combined with the discovery of transitions between (highly correlated) states in which the grid is the same but the peak firing rate of each field is different [20], and with a new understanding of the disorder and frustration inherently associated to the grid representation of curved environment [32], puts to rest the rigid order which had appeared as the most salient character of the newly discovered grid cells. It suggests instead a sort of spin glass at intermediate temperature, i.e., that in order to code densely and efficiently for position on (many) continuous manifolds, grid cells have to be equipped with the flexibility and the ability to compromise characteristic of self-organized disordered system.

## ACKNOWLEDGMENTS

Work supported by Human Frontier grant RGP0057/2016.

## Appendix A: Mean field equations: Binary Model

The free-energy can be written, in the large *N* limit, in terms of macroscopic quantities:

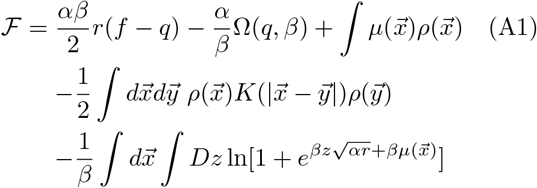

where *β* is an inverse temperature or noise level, and the function Ω(*q, β*) is given by

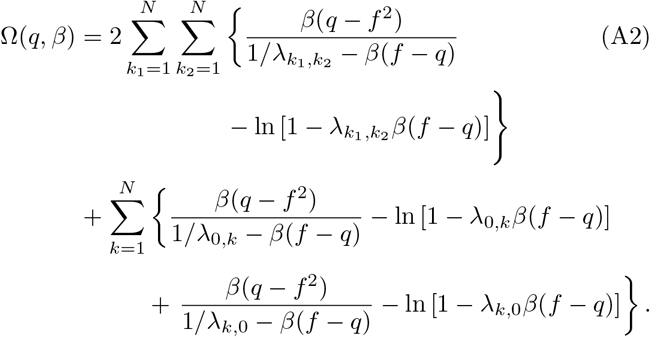

The order parameters minimizing the free energy functional are the average activity 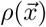 (see main text) and

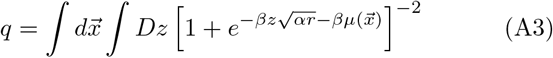

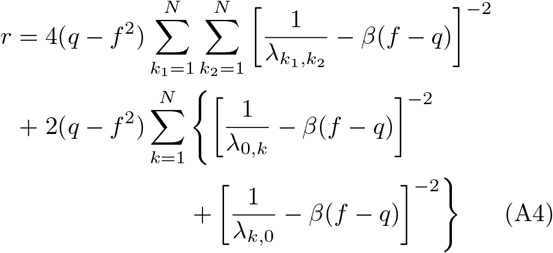

where *λ* enforces the constraint 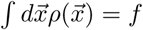 and 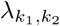 are the eigenvalues of the kernel *K* and are given by

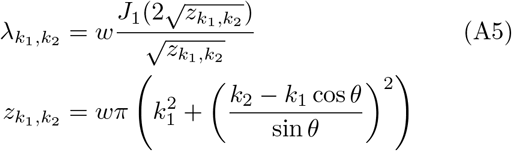

where *J*_1_ is the Bessel function of the first kind of order 1.

In the text we focus on the limit of vanishing stochastic noise *β → ∞*, and the term *β*(*q −f*), which remains finite in such limit, can be identified with the parameter *ψ* of the threshold-linear model, quantifying the reverberation through the loops of the network of the *quenched* noise, which is due to the interference of the other maps.

## Appendix B: Mean field equations: Threshold-linear Model

When an energy functions can be defined (with full or in any case symmetric connectivity) the thermodynamics of the system is dominated by the minima of the free energy *density*

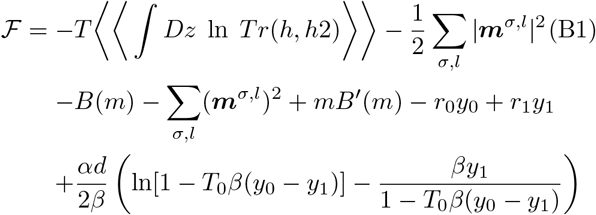

where we have maintained a notation consistent with [31] and [24], for example

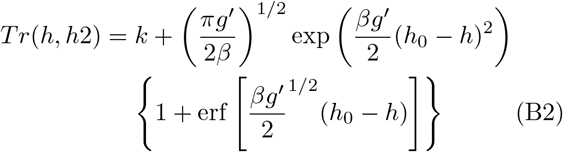

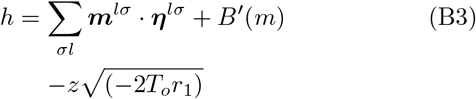

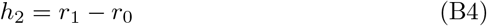

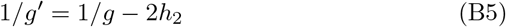

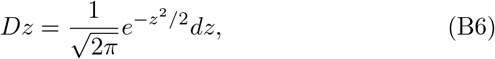

while 〈〈⋅〉〉 denotes an average over the quenched noise (the field centers in all other stored maps, distinct from the one which is currently expressed); and *B*(*x*), together with the gain *g*, can be used to constrain the mean activity and the sparsity of the activity pattern [31], analogous to the parameter *λ* in the binary model.

The minima are given, in the limit *T* → 0, by the saddle point equations

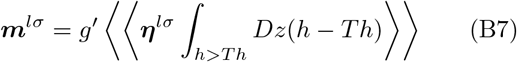

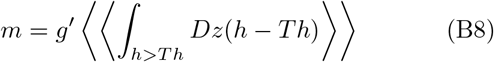

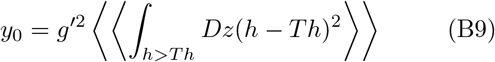

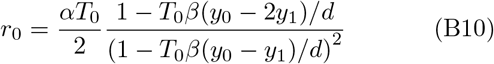

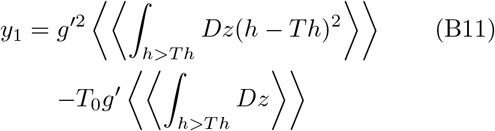

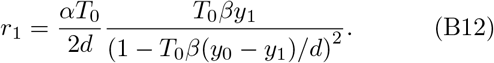

Introducing the variables

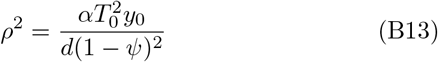

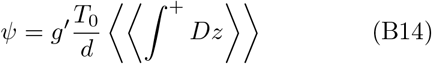

we can write the free energy as a function of macroscopic
quantities

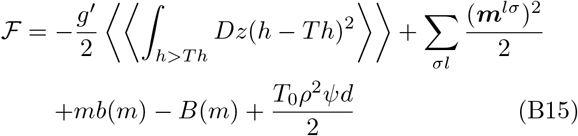

with now

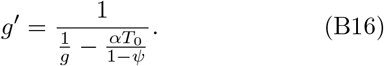

To calculate the storage capacity, we focus on the case in which a single environment is retrieved by the network,

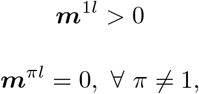

although the analysis can be extended to the retrieval of bump states that are localized in multiple environments. Without loss of generality, we assume therefore that environment *π* = 1 is retrieved. With this assumption, and introducing the two signal-to-noise ratios

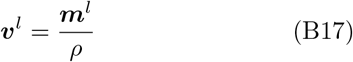

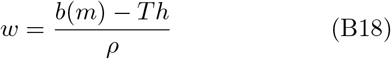

that represent respectively the *environment specific* component of the signal and the *uniform* background inhibition acting on each unit, the saddle point equations can then be reduced to a system of two equations in two variables

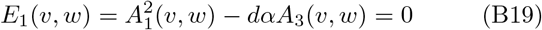

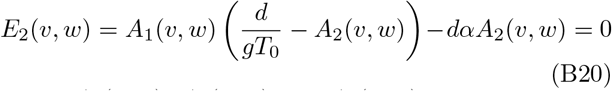

where *A*_1_(*w*; *v*), *A*_2_(*w*; *v*) and *A*_3_(*w*; *v*) are the averages:

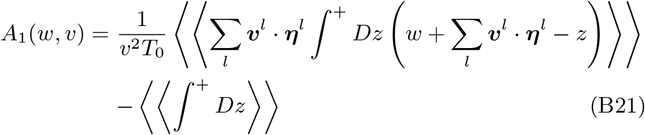

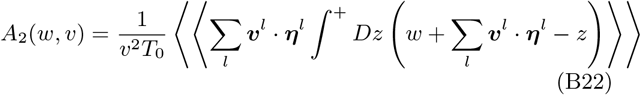

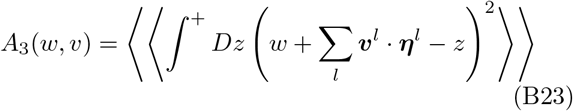

Solutions to equations (B19) and (B20) give the minima of the free energy that correspond to the retrieval of one of the stored environments. *E*_1_(*v, w*) = 0 describes a closed curve in the *w − v* plane, and these solutions are the intersections with *E*_2_(*v, w*) = 0, which depends on the gain *g*.

As the storage load *α* = *p/C* increases, this closed curve shrinks and eventually disappears. The value *α* = *α*_*c*_ at which the curve vanishes marks a phase transition: for *α > α*_*c*_ retrieval solutions do not exist. The storage capacity *α*_*c*_ can therefore be calculated by finding the vanishing point of *E*_1_ = 0, and in this way one automatically selects the optimal value of the gain *g*, which therefore

## Appendix C: Finite connectivity and noise reverberation

Equations B19 and B20 can be extended to arbitrary value of connectivity density *C/N* following the self-consistent signal-to-noise analysis developed in [30]. This gives

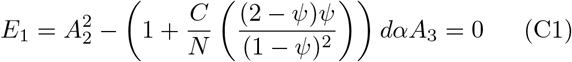

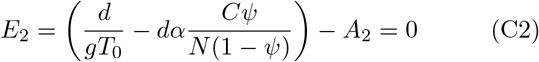

These equations interpolate, as the free parameter *C/N* varies, between the two limiting cases of a fully connected network (*C/N* = 1) and the extremely diluted case (*C/N* → 0) studied in [36]. We see that the reverberation factor *ψ* enters in the equation for the storage capacity as a correction on the loopless equation 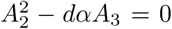, modulated by the connectivity density *C/N*, and that the lower the *ψ*, the higher the storage capacity.

For the fully connected network this correction gives

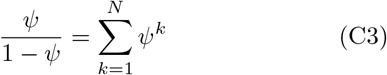

which is the sum over all the k-loops contributions to the reverberation of the noise.

Note, finally, that for ease of comparison with the binary model we have written in the main text

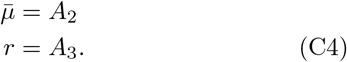

## Appendix D: Free-energy barriers in the binary model

Free-energy values for the different metastable states are calculated using (A4) after order parameters have been computed by solving the saddle-point equations. These equations are solved iteratively, starting from an initial condition for order parameters, and iterating the values of the order parameters until convergence to fixed values. The free-energiy values of the different metastable states are obtained by initializing 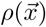 as 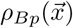 for Bump States (Fig.6(a,e)) and 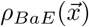 for Band Edge states (Fig.6(b,f)) or 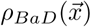 for Band Diagonal states (Fig.6(c,g)). In order to estimate the size of the barrier that must be jumped over in order to go from one state *X* to another state *Y*, we proceed as follows. The activity profile is initialized as 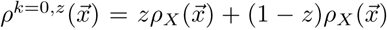, with z chosen such that 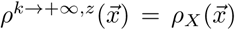 and 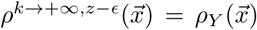 for *ϵ ≪ z*. When solving equations from such an initial condition, the network state goes close to a saddle-point lying at the boundary between the two basins of attraction associated to states *X* and *Y*, before sliding into state *X* as shown in Fig.7(c). The size of the barrier is then given by the difference between the free-energy of the saddle-point and that of the meta-stable state *X*.

